# A bacteriophage split DNA polymerase incorporates a sliding clamp-derived processivity subunit

**DOI:** 10.64898/2026.09.30.755723

**Authors:** Dwaipayan Basu, Ryan K. Min, Koe Inlow, Joe Pogliano, Elizabeth Villa, Kevin D. Corbett

## Abstract

Viruses that infect bacteria, termed bacteriophages or phages, typically encode their own DNA replication machinery that rapidly generates many copies of the phage genomic DNA in an infected cell. Nucleus-forming phages of the chimallivirus family possess a distinctive repertoire of predicted DNA replication proteins, with two proteins annotated as DNA polymerases plus a DnaB-like replicative helicase, but lacking an identified primase or sliding clamp protein. Here, we demonstrate that in the *Pseudomonas aeruginosa*-infecting chimallivirus PhiKZ, the core DNA polymerase comprises a stable 1:1 complex of two DNA polymerase subunits, termed DNAP^N^ and DNAP^C^. A 2.1 Å resolution cryoelectron microscopy structure of this core complex reveals homology to other phage and bacterial B-family DNA polymerases. Next, we identify a direct but transient interaction between the core DNA polymerase complex and a third protein termed Rhc, which possesses an N-terminal RNase H domain and a C-terminal domain related to DNA sliding clamp proteins. Cryoelectron microscopy structures of a stabilized DNA polymerase-Rhc complex, and of this complex bound to a primer-template DNA, reveal a single Rhc subunit that cooperates with the DNA polymerase thumb domain to bind double-stranded DNA emerging from the DNA polymerase active site. While the Rhc RNase H domain is not resolved in our structures, this domain is situated such that it could degrade RNA associated with an incoming template DNA strand. *In vitro* primer-extension assays show that Rhc strongly boosts DNA polymerase activity, and that this boost relies on the Rhc-DNA polymerase interface we observe in our structures. Overall, our findings reveal a distinctive chimallivirus-specific DNA replication machine with an integrated DNA-binding subunit related to sliding clamp proteins, and a flexibly-linked RNase H domain.

## INTRODUCTION

Bacteriophages (phages) are the most numerous organisms on the planet, and show high diversity in their mechanisms of infection, replication, and evasion of host immune responses. While historical isolation and cultivation methods have selected for phages with genomes less than ∼175 kb, recent work has demonstrated that “jumbo” phages with genomes over 200 kb are widespread in nature^1^. One family of jumbo phages is termed *Chimalliviridae* (chimalliviruses) after a characteristic protein, Chimallin/ChmA (also termed PhuN) which forms a protective shell around the replicating phage genome in an infected host^2–4^. This “phage nucleus”, which is preceded in the viral life cycle by the membrane-bound early phage infection (EPI) vesicle, protects the phage genome from direct attack by host-encoded restriction endonucleases, CRISPR-Cas nucleases, and other DNA-targeting immune pathways^5–9^. At the same time, the phage nucleus introduces challenges for the phage, including the need to transport mRNAs produced in the phage nucleus to the cytoplasm for translation, and the need to import specific phage and host proteins into the phage nucleus to support DNA replication and transcription.

Chimalliviruses show a naturally broad host range^10^, making them promising candidates for phage therapy: the use of phages to combat life-threatening antibiotic-resistant infections in human patients^11,12^. There is considerable interest in genome engineering of chimalliviruses to further improve their utility in phage therapy, for example by integrating proteins that expand host range and/or counter specific host immune pathways for which these phages lack natural defenses^13^. These efforts are hampered by a lack of knowledge of chimalliviruses’ replication mechanisms: while the phage nucleus and EPI vesicle are well-characterized, key questions such as how DNA is replicated and packaged into new virions in this phage family remain unknown. A recent bioinformatics study of diverse jumbo phages showed that chimalliviruses possess a distinctive set of DNA replication and recombination proteins^14^. While most phages encode a single DNA polymerase with an integrated 3′-5′ exonuclease domain, chimalliviruses encode separate proteins annotated as putative DNA polymerase and exonuclease proteins. Chimalliviruses also encode a divergent DnaB-type replicative helicase, but other canonical DNA replication proteins including primase, sliding clamp, and clamp loader have not been identified. These features indicate that chimalliviruses possess a distinctive and highly diverged set of genome replication machinery.

Here, we reconstitute the DNA polymerase of the *Pseudomonas aeruginosa*-infecting chimallivirus PhiKZ, and show that it comprises three proteins: the previously identified DNA polymerase and exonuclease subunits form a stable core that transiently associates with a third protein termed Rhc (<u>R</u>Nase <u>H</u>-<u>c</u>lamp). Rhc possesses an N-terminal RNase H domain and a C-terminal domain related to DNA sliding clamp proteins. Using cryoelectron microscopy, we determine high-resolution structures of the three-subunit PhiKZ DNA polymerase and its complex with DNA, revealing overall similarity to known DNA polymerases and an expanded DNA binding interface on Rhc’s clamp domain. Overall, this work defines the architecture and mechanism of a multi-subunit bacteriophage DNA polymerase with a distinctive bifunctional DNA-binding subunit that also possesses an RNase H domain.

## RESULTS

### Structure of the PhiKZ DNAP^N+C^ complex

To determine the structural basis for DNA replication in chimalliviruses, we expressed and purified a 1:1 complex of DNAP^N^ (gp82) and DNAP^C^ (gp50) from the *Pseudomonas* phage PhiKZ (**Figure S1A**), and determined a 2.14 Å resolution cryoEM structure of this complex (**Figure S1B-G**). The structure reveals an overall architecture similar to B-family polymerases, with N-terminal domain (NTD), exonuclease, palm, fingers, and thumb domains (**Figure 1A-C**). The subunit split occurs within a large insertion in the palm domain (discussed below), such that DNAP^N^ possesses the NTD, exonuclease, and part of the palm domain, and DNAP^C^ possesses the remainder of the palm domain plus fingers and thumb domains (**Figure 1A-C**). Thus, despite being encoded in two genes, the PhiKZ DNAP^N+C^ complex is a stable polymerase/exonuclease complex like other B-family DNA polymerases.

**Figure 1.**
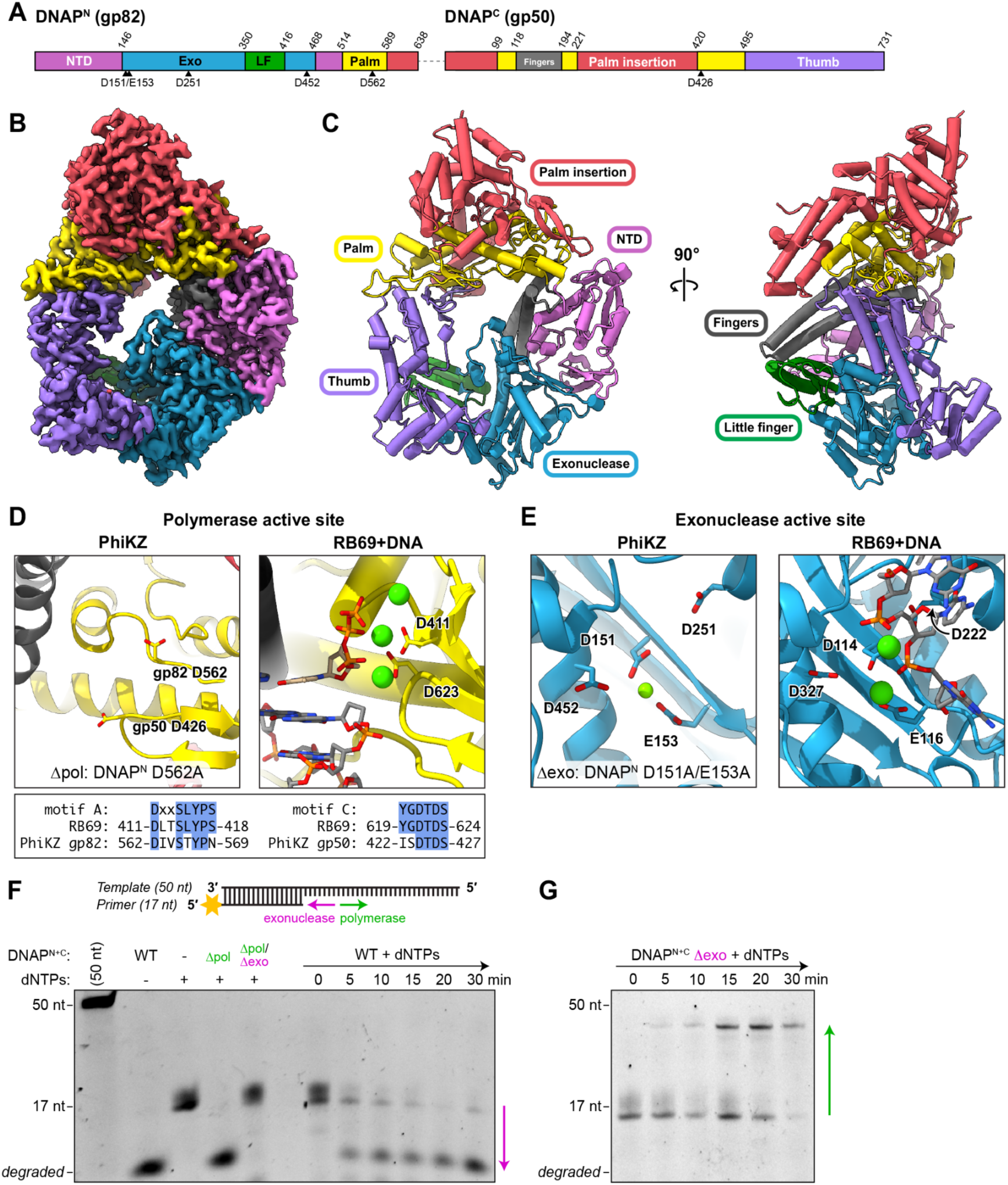
Structure and activity of PhiKZ DNAP^N+C^. (A) *Top:* Domain diagram of PhiKZ DNAP^N+C^ (DNAP^N^ (gp82) above, DNAP^C^ (gp50) below), with domains colored and labeled. (B) View of the 2.14 Å resolution cryoEM map of PhiKZ DNAP^N+C^ (see **Figure S1** for structure determination details), with domains colored as in panel (A). (C) Two views of the PhiKZ DNAP^N+C^ complex, with domains colored as in panel (A) and labeled. (see **Figure S2** for comparison with RB69 DNA polymerase structure) (D) Closeup views of the polymerase active site of PhiKZ DNAP (left) and RB69 (right; PDB ID 1IG9)^16^, with domains colored as in panel (A) and conserved active site residues shown in sticks. Green spheres indicate Ca^2+^ ions bound in the RB69 structure. *Bottom:* Alignment of polymerase motifs A and C in RB69 and PhiKZ DNAP^N+C^ (ref. ^18^). (E) Closeup views of the exonuclease active site of PhiKZ DNAP (left) and the editing complex of RB69 DNAP (right; PDB ID 1CLQ)^17^, with domains colored as in panel (A) and conserved active site residues shown in sticks. Green spheres indicate bound Mg^2+^ (for PhiKZ) or Ca^2+^ (for RB69) ions. (F) *Top:* Schematic of primer extension assay with 50-nt template strand and 17-nt labeled primer. See **Materials and Methods** for sequences. *Bottom:* Primer extension assay with PhiKZ DNAP^N+C^ and the indicated DNA substrate. DNAP^N+C^ indicates the two-subunit gp82-gp50 complex (coexpressed and purified), Δpol indicates polymerase-dead enzyme (DNAP^N^/gp82 D562A), and Δexo indicates exonuclease-dead enzyme (DNAP^N^/gp82 D151A/E153A). Δpol/Δexo indicates a double-catalytic dead enzyme (DNAP^N^/gp82 D151A/E153A/D562A). All assays were performed with 25 nM annealed primer-template DNA and 300 nM enzyme. (G) Primer extension assay with DNAP^N+C^ Δexo (DNAP^N^/gp82 D151A/E153A).

The PhiKZ DNAP^N+C^ structure is most similar to that of the *Escherichia* phage RB69 DNA polymerase, a single-subunit B-family polymerase whose structure and function has been extensively studied. Structures of the RB69 DNAP have been determined in the apo state^15^ and bound to DNA in states mimicking strand extension/polymerization^16^ and editing/degradation^17^. Overall, our structure of PhiKZ DNAP^N+C^ in the apo state most closely resembles the “open” apo state of the RB69 DNAP, with the fingers domain extended away from the palm domain (**Figure 1A-C, Figure S2**). In each individual domain, the PhiKZ DNAP^N+C^ structure is highly similar to the RB69 DNAP structure despite relatively low overall sequence identity of 28.8%, spanning the NTD (6.4 Å r.m.s.d. over 96 Cɑ atoms), exonuclease (4.6 Å r.m.s.d. over 176 Cɑ atoms), palm (4.5 Å r.m.s.d. over 78 Cɑ atoms), fingers (3.3 Å r.m.s.d. over 72 Cɑ atoms), and thumb (5.8 Å r.m.s.d. over 160 Cɑ atoms) domains (**Figure S2**).

Both the polymerase and exonuclease active sites are highly conserved with RB69 DNAP and other B-family polymerases. The DNA polymerase active site contains two absolutely conserved aspartate residues, D562 in DNAP^N^ (within “Motif A”) and D426 in DNAP^C^ (within “Motif C”)^18^ (**Figure 1D**). The exonuclease active site contains four negatively-charged residues in DNAP^N^ (D151, E153, D251, and D452) which coordinate one Mg^2+^ ion in the DNAP^N+C^ structure (**Figure 1E**). Based on these data and prior work on B-family DNA polymerases, we designed a Δpol mutant (DNAP^N^ D562A) and a Δexo mutant (DNAP^N^ D151A/E153A) for biochemical studies.

To directly measure DNA polymerase activity of the DNAP^N+C^ complex, we developed an *in vitro* polymerase assay using a 17-nucleotide labeled primer and a 50-nucleotide template strand (**Figure 1F**). The wild-type DNAP^N+C^ complex degraded the labeled primer, demonstrating that its exonuclease activity predominates in solution over its polymerase activity (**Figure 1F**). In contrast, the Δexo mutant extended the primer to the full length of the unlabeled template strand (**Figure 1G**). A double Δpol/Δexo mutant showed neither polymerization nor degradation (**Figure 1F**), demonstrating that the observed activities are specific to the purified DNAP^N+C^ complex.

The PhiKZ DNAP is significantly larger (DNAP^N^: 638 residues; DNAP^C^: 731 residues; total: 1369 residues) than RB69 DNAP (903 residues). The additional size comes mainly from two large insertions, which we term the “palm insertion” domain and the “little finger” domain. The palm insertion domain is a largely ɑ-helical domain comprising three segments: DNAP^N^ residues 589-638, DNAP^C^ residues 1-98, and DNAP^C^ residues 221-419 (**Figure 1A**). These segments fold together into a single domain that encloses nearly the entire palm domain (**Figure 1C**). Structure-similarity searches with DALI^19^ and Foldseek^20^ do not reveal any structural relatives of this domain. Sequence alignments and structure predictions indicate that the palm insertion domain is highly conserved across chimallivirus DNAPs.

The second insertion in the PhiKZ DNAP, which we term the little finger (LF) domain, is inserted into the exonuclease domain and spans DNAP^N^ residues 350-415. The LF domain adopts a mixed ɑ-β fold, and the DALI structure comparison server^19^ indicates that it is similar to the little finger (LF) or polymerase associated domain (PAD) found in Y-family DNA polymerases (**Figure S3A-B**). In Y-family DNAPs, the LF domain is fused to the C-terminus of the thumb domain and associates with the major groove of the double-stranded primer-template DNA, with two β-strands forming hydrogen bonds with backbone phosphates along both DNA strands^21–23^. The LF domain strengthens DNA binding, enhances processivity, and plays a role in Y-family DNAPs’ ability to bypass DNA lesions^24,25^. While the PhiKZ DNAP LF domain shares a common fold with Y-family DNA polymerases’ LF domains, it is inserted into the exonuclease domain rather than fused to the thumb domain, and is positioned behind the palm domain where it is unlikely to interact with DNA. Moreover, sequence alignments of chimallivirus DNAP^N^ subunits show that the LF domain is found only in a subset of chimalliviruses that infect *Pseudomonas* (**Figure S3C**). These observations suggest that the PhiKZ DNAP LF domain may be involved in protein-protein interactions, perhaps with another protein found only in this subset of phages, rather than DNA interactions as in Y-family DNAPs. In *in vitro* primer extension assays, deletion of the LF domain has a minor effect on the DNA polymerase activity of the DNAP^N+C^ complex (**Figure S3D**).

### A bifunctional RNase H-clamp protein contributes to DNAP^N+C^ activity

Concurrent with this work, a concurrent study of the chimallivirus RNase H-like protein Rhc (<u>R</u>Nase <u>H</u>-<u>c</u>lamp) identified a direct interaction between this protein and DNAP^C^, first by proteome-wide AlphaFold screening and later by immunoprecipitation and mass spectrometry from phage-infected cells (Rani et al. *bioRxiv*). Separately, antisense oligonucleotide–mediated knockdown of PhiKZ Rhc (gp155) was shown to compromise phage DNA replication in infected cells^26^. We confirmed the likelihood of a direct DNAP-Rhc interaction using AlphaFold 3, which showed a pairwise interface predicted template modeling (ipTM) score of 0.70 between DNAP^C^ and Rhc (**Figure 2A-B**). An AlphaFold 3 prediction incorporating all three proteins – DNAP^N^, DNAP^C^, and Rhc – showed an overall ipTM of 0.80, in the range associated with highly confident predictions of complex formation^27^. With the addition of primer-template DNA and Mg^2+^ ions, the overall AlphaFold 3 ipTM rises to 0.90 (**Figure S4A-B**).

**Figure 2.**
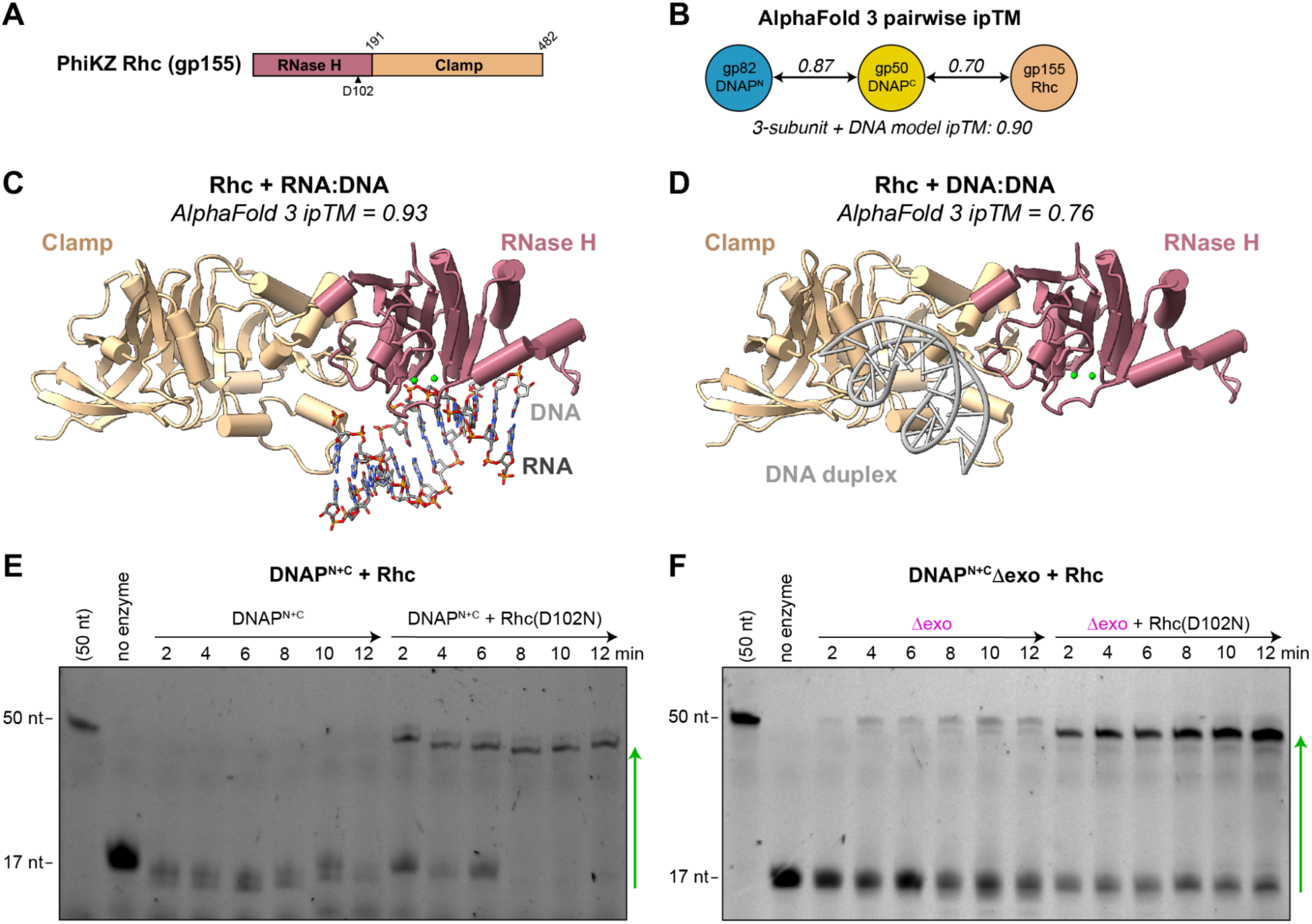
PhiKZ Rhc is a predicted RNase H-clamp fusion that contributes to DNAP^N+C^ catalysis. (A) Domain schematic of PhiKZ Rhc (gp155). (B) AlphaFold 3 pairwise ipTM scores for PhiKZ DNAP^N^, DNAP^C^, and Rhc. See **Figure S4** for AlphaFold 3 prediction of the full 3-subunit model with primer-template DNA. (C) AlphaFold 3 prediction of PhiKZ Rhc bound to an RNA:DNA heteroduplex. See **Figure S5** for details. (D) AlphaFold 3 prediction of PhiKZ Rhc bound to a DNA:DNA duplex. See **Figure S6** for details. (E) Primer extension assay with PhiKZ DNAP^N+C^, in the absence and presence of equimolar Rhc (D102N catalytic-dead mutant). (F) Primer extension assay with PhiKZ DNAP^N+C^ Δexo (DNAP^N^/gp82 D151A/E153A), in the absence and presence of equimolar Rhc (D102N catalytic-dead mutant).

AlphaFold models of PhiKZ Rhc predict that the protein has two domains: an N-terminal RNase H1 domain homologous to RnhA from *E. coli* and other bacteria (**Figure 2C, Figure S5**); and a C-terminal domain distantly related to DNA sliding clamp proteins including eukaryotic PCNA (proliferating cell nuclear antigen) and the 9-1-1 complex (RAD9, RAD1, HUS1) (**Figure 2D, Figure S6, Table S2**). Close comparison of the Rhc C-terminal domain with sliding clamp proteins reveals that Rhc lacks the structural elements responsible for these proteins’ oligomerization into DNA-encircling rings (**Figure S6D**); in keeping with this observation, AlphaFold was unable to confidently predict a structure for an Rhc oligomer (not shown). An AlphaFold prediction of Rhc with an RNA:DNA heteroduplex generated a confident model with the predicted RNase H active site poised for cleavage of the RNA strand (**Figure 2C, Figure S5**). In contrast, an AlphaFold prediction of Rhc with a double-stranded DNA generated a confident model with the DNA bound to a large, concave, positively charged surface on the Rhc clamp domain (**Figure 2D, Figure S6**). Superimposing this model onto structures of sliding clamp trimers shows that this surface corresponds to the inner, DNA-associated surface of these sliding clamp proteins (**Figure S6D**).

To test whether Rhc functionally collaborates with DNAP^N+C^, we tested the effect of adding Rhc to our *in vitro* primer extension assays. We were unable to express and purify wild-type PhiKZ Rhc in *E. coli* due to toxicity, but we could purify an RNase H catalytic-dead mutant protein (D102N). Strikingly, addition of Rhc(D102N) converted DNAP^N+C^ catalytic activity from predominantly degrading the primer, to robustly extending it (**Figure 2E**). Similarly, addition of Rhc to the DNAP^N+C^ Δexo mutant resulted in a dramatic increase in the enzyme’s DNA polymerization activity (**Figure 2F**). These data show that Rhc strongly contributes to the DNA polymerase activity of DNAP^N+C^, independently of its RNase H catalytic activity.

### Structure of DNAP^N+C^ bound to Rhc

We next sought to reconstitute a complex of DNAP^N^, DNAP^C^, and Rhc for structure determination. We expressed and purified PhiKZ Rhc(D102N), mixed it with the purified DNAP^N+C^ complex, and prepared samples for cryoEM analysis. An initial cryoEM structure using mixed DNAP^N+C^ and Rhc showed only density for the DNAP^N+C^ complex, suggesting that Rhc does not stably associate with DNAP^N+C^. Because the C-termini of Rhc and DNAP^C^ were predicted by AlphaFold to be closely juxtaposed in the complex, we fused these two proteins to designed peptides (termed SYNZIPs) that were previously shown to form specific coiled-coil heterodimers (**Figure S4C-D**)^28^. Separately-expressed DNAP^N^+DNAP^C^-SYNZIP1 and Rhc(D102N)-SYNZIP2 proteins partially co-eluted on a size exclusion column (**Figure S7A-B**), and we could determine a 2.8 Å resolution cryoEM structure of this complex that revealed strong density for DNAP^N+C^ plus the clamp domain of Rhc (**Figure 3A-C, S7C-G**). In the cryoEM structure, the Rhc RNase H domain is not visible, presumably due to flexibility; local refinement was unable to recover density for this domain. We also do not observe density for the SYNZIP1/2 coiled-coil segments.

**Figure 3.**
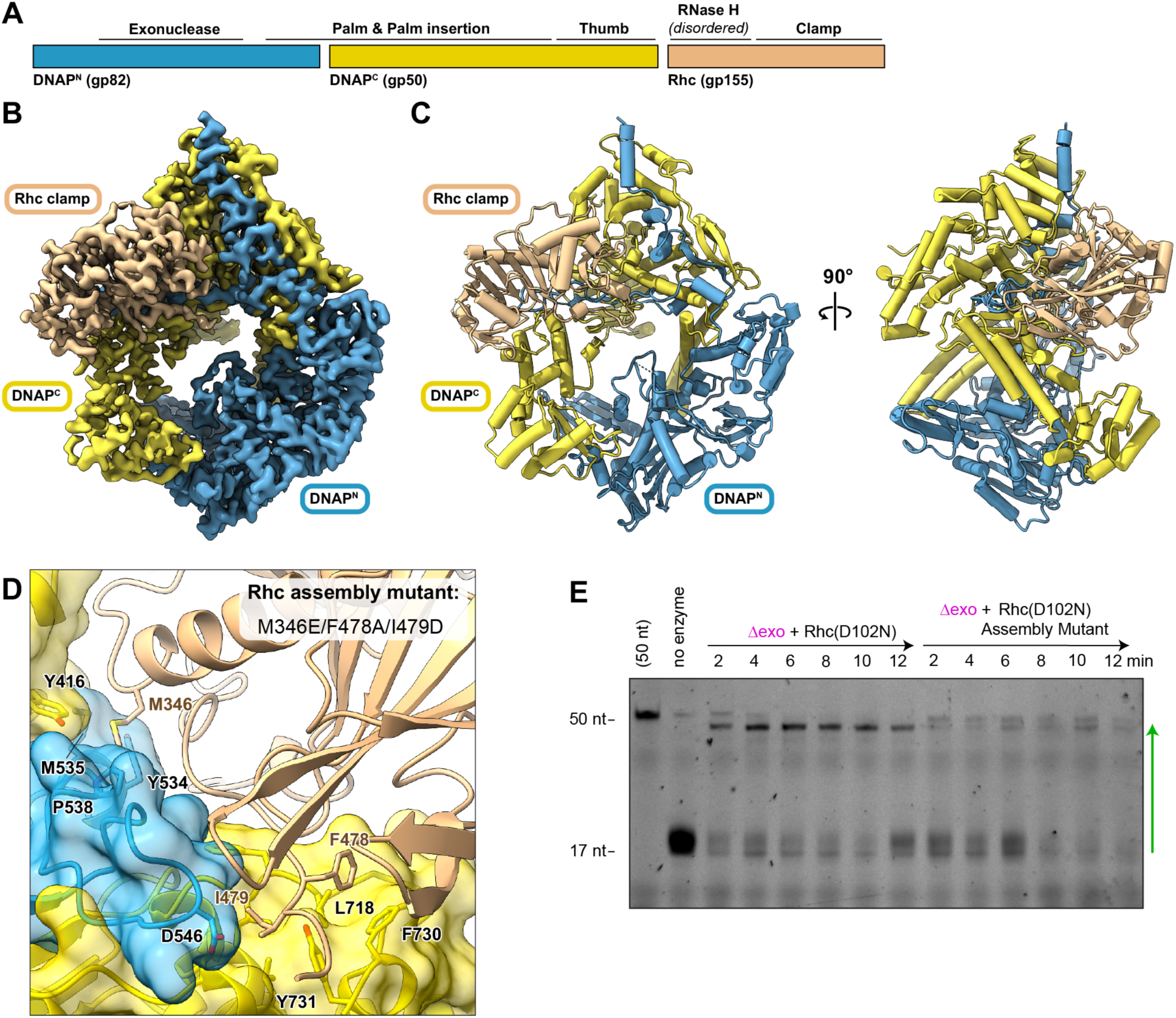
Structure of the PhiKZ DNAP^N+C^-Rhc complex. (A) Subunit schematic of the three-subunit PhiKZ DNAP^N+C^-Rhc complex (B) CryoEM density at 2.82 Å resolution for the three-subunit PhiKZ DNAP^N+C^-Rhc complex, comprising DNAP^N^ (gp82; blue), DNAP^C^ (gp50; yellow), and Rhc (gp155; light brown). The Rhc RNase H domain is disordered and not visible. See **Figure S7** for structure determination details. (C) Two views of the PhiKZ DNAP^N+C^-Rhc complex. The Rhc RNase H domain is disordered and is not observed in the structure. (D) Closeup of the interface between Rhc and the DNAP^N+C^ complex, with three Rhc residues mutated in the assembly mutant (M346, F478, and I479) shown as sticks, along with their interacting residues on the DNAP^N^ and DNAP^C^ subunits. (E) Primer extension assay with PhiKZ DNAP^N+C^ Δexo (DNAP^N^/gp82 D151A/E153A), in the presence of equimolar Rhc (D102N catalytic-dead mutant) or the Rhc (D102N) assembly mutant (M346E/F478A/I479D).

In the structure of DNAP^N+C^ bound to Rhc(D102N), the Rhc clamp domain forms an extensive interface with the thumb domain of DNAP^C^, plus the palm and palm insertion domains composed of both DNAP^N^ and DNAP^C^, burying a total of ∼1,800 Å^2^ on Rhc (**Figure 3C-D**). Unlike the extensively-interdigitated and mostly hydrophobic interaction between DNAP^C^ and DNAP^N^, the interface between Rhc and DNAP^N+C^ includes a mix of hydrophobic and polar/charged residues on both partners.

Based on the structure, we designed an Rhc “assembly mutant” to disrupt the Rhc-DNAP interface (M346E/F478A/I479D). The Rhc(D102N) assembly mutant expressed and purified equivalently to Rhc(D102N), but did not boost DNA polymerization activity of DNAP^N+C^ *in vitro* (**Figure 3E**). This experiment supports the validity of the observed Rhc-DNAP interface and the protein’s role in boosting DNA polymerase activity.

In the structure of DNAP^N+C^ bound to Rhc, the Rhc RNase H domain is not visible, presumably because of flexibility between the RNase H and clamp domains. In an AlphaFold 3 prediction of the PhiKZ DNAP^N+C^– Rhc complex structure bound to primer-template DNA (**Figure S4**), the overall architecture of the three subunits closely matches our cryoEM structure. In this prediction, the RNase H domain of Rhc is poised above the template DNA strand, such that its active site would be properly positioned to degrade an RNA strand annealed to the incoming template strand prior to replication. This positioning suggests that, in addition to providing a large DNA binding surface through its clamp domain, Rhc could degrade an RNA strand annealed to the template DNA, thereby promoting replication.

### Structure of a DNA-bound DNAP^N+C^–Rhc complex

We next attempted to visualize a DNA-bound structure of the DNAP^N+C^–Rhc complex. We annealed a 50-nucleotide template strand to a 16-nucleotide primer with a 3′ dideoxycytosine to prevent strand extension, and incubated the purified DNAP^N+C^–Rhc complex with a 5x molar excess of this DNA substrate plus dCTP as an incoming base. We determined a 2.76 Å resolution structure of the resulting complex by cryoEM (**Figure 4A-B, Figure S8**). The structure reveals a 16-base pair double-stranded DNA segment representing the annealed primer-template DNA complex, but no further density extending into either the DNA polymerase or exonuclease active sites. The primer-template DNA is bound between the DNAP^C^ thumb domain and the Rhc clamp domain, both of which bind DNA through large, positively-charged surfaces (**Figure 4C-D**). This structure confirms that Rhc’s clamp domain contributes to binding replicated double-stranded DNA emerging from the DNAP active site, thereby likely boosting the enzyme’s overall affinity for DNA and its processivity.

**Figure 4.**
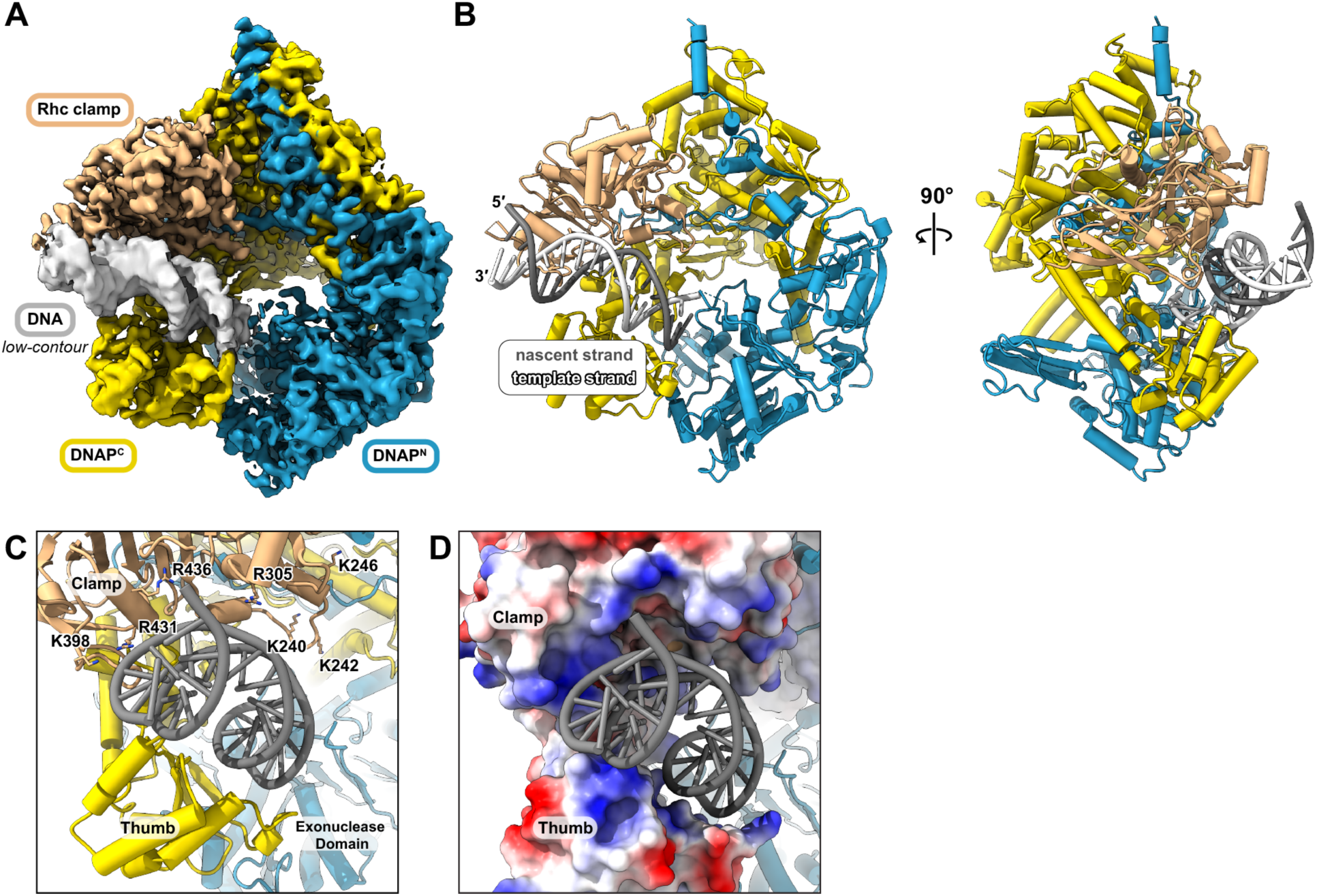
Structure of the PhiKZ DNAP^N+C^-Rhc + DNA complex. (A) CryoEM density at 2.76 Å resolution for the three-subunit PhiKZ DNAP^N+C^-Rhc complex (subunits colored as in Figure 3) bound to DNA (light gray, contoured at a lower level for easier visibility). See **Figure S8** for structure determination details. (B) Two views of the PhiKZ DNAP^N+C^-Rhc + DNA complex. The template and primer strands are colored white and dark gray, respectively. No density is observed for the template or primer strands in the DNAP^N+C^ active sites. (C) Closeup view of the double-stranded primer-template DNA bound between the Rhc clamp and the DNAP thumb domains. Positively-charged residues on the interacting surface of Rhc are shown as sticks and labeled. (D) View as in panel (C), showing an electrostatic surface for the Rhc clamp domain and the DNAP thumb domain.

## DISCUSSION

Chimalliviruses possess a distinctive complement of predicted DNA replication and repair proteins, with two proteins showing homology to B-family DNA polymerases plus a DnaB-like replicative helicase, but conspicuously lacking primase or sliding clamp proteins^14^. Here we show that PhiKZ’s two predicted DNA polymerase proteins form an extensively-interdigitated 1:1 heterodimer with an overall architecture similar to other B-family DNA polymerases, including that of phage RB69. *In vitro*, the PhiKZ DNAP^N+C^ complex shows both polymerase and exonuclease activity.

Recently, antisense oligonucleotide–mediated knockdown of a predicted RNase H protein in PhiKZ (gp155; here termed Rhc) was shown to compromise phage DNA replication in infected cells^26^. Curiously, expression of a catalytic-dead mutant of Rhc in these cells was sufficient to fully rescue DNA replication and phage titer, demonstrating that Rhc has a non-catalytic role in DNA replication^26^. Here, we explain this observation by demonstrating that PhiKZ Rhc associates directly with the DNAP^N+C^ complex and boosts its polymerase activity *in vitro*. Our cryoEM structure of DNAP^N+C^ bound to Rhc and DNA shows that Rhc’s C-terminal clamp domain is structurally related to DNA sliding clamps, but that rather than encircling DNA, this domain provides a large binding surface for the replicated DNA emerging from the polymerase active site. In this sense, Rhc is similar to sliding clamp-derived processivity factors in several eukaryotic viruses, especially the UL42 protein in herpes simplex virus (HSV-1) which associates with the DNA polymerase and aids binding to replicated double-stranded DNA as a monomer^29–32^. Unlike herpesvirus UL42, Rhc does not stably associate with the DNAP^N+C^ complex *in vitro*, meaning that it is likely to only associate with DNAP^N+C^ when the polymerase is bound to a template DNA and polymerizing a complementary strand. While the Rhc RNase H domain is disordered in our structure, its inferred positioning suggests that it could degrade an RNA strand annealed to the template DNA strand.

Chimalliviruses are distinctive in that they enclose their replicating genomic DNA inside a “phage nucleus”, a protein-bounded compartment that excludes host factors including ribosomes^3,33^. This segregation requires that mRNAs, which are transcribed within the phage nucleus, be transported out of this compartment for translation. Typically, ribosome binding and translation of bacterial and phage mRNAs reduces the propensity for these mRNAs to re-anneal to the template DNA strand to form R-loops. The physical segregation of transcription from translation in chimalliviruses may increase the propensity for these phages’ mRNAs to form R-loops. In this scenario, an RNase H associated with the phage DNA polymerase may contribute to DNA replication by degrading R-loops ahead of the enzyme. While in PhiKZ, DNA replication of an Rhc knockdown is fully complemented by a catalytic-dead mutant^26^, a concurrent study in the *E. coli* phage Goslar shows that in this phage, Rhc plays both non-catalytic and catalytic roles in DNA replication: a catalytic-dead Rhc partially complements the DNA replication defect observed upon CRISPR-mediated Rhc knockdown, while wild-type Rhc fully complements this defect (Rani et al. *bioRxiv*). Thus, while different chimalliviruses share a dependence on Rhc’s clamp domain to promote DNA replication, they vary in their requirement for Rhc’s RNase H activity.

The observation that Rhc only transiently associates with DNAP^N+C^ means that the protein could have other roles in the phage replication cycle. To that end, a concurrent study shows that knockdown of Goslar Rhc strongly affects transcription of a subset of late genes; that this activity depends on both Rhc’s clamp domain and its RNase H catalytic activity; and that this activity is independent of DNA replication (Rani et al. *bioRxiv*). This observation suggests a role for Rhc in resolution of R-loops, which (as described above) may be prevalent in chimalliviruses because of the physical segregation of transcription from translation, and may strongly compromise transcription of particular genes if left unresolved.

### Limitations of the study

While we show that Rhc contributes to the DNA polymerase activity of DNAP^N+C^ *in vitro* and directly binds replicated DNA, our assays do not reveal whether Rhc’s specifically increases processivity or DNA replication rate of DNAP^N+C^. Our structure suggests that Rhc stabilizes DNA binding of DNAP^N+C^, suggesting that it increases processivity; proving this point will require more quantitative assays.

Our *in vitro* primer-extension assays used an Rhc catalytic-dead mutant (D102N), because we were unable to express and purify the wild-type protein from *E. coli*. Thus, we are only able to measure a non-catalytic role for Rhc in DNA replication *in vitro*. A prior study used cell-free expression to produce small quantities of both wild-type PhiKZ Rhc and the catalytic-dead D102N mutant^26^. This study showed that wild-type Rhc efficiently degrades RNA:DNA hybrids *in vitro*, and that the D102N mutant is inactive^26^.

## MATERIALS AND METHODS

### Protein expression and purification

The coding sequences for PhiKZ gp82 (NCBI # NP_803648.1) and gp155 (NCBI # NP_803721.1) were cloned into UC Berkeley Macrolab vector 2-BT (Addgene #29666), which encodes an N-terminal TEV-protease-cleavable His_6_ tag. The coding sequence for PhiKZ gp50 (NCBI # NP_803616.1) was cloned into UC Berkeley Macrolab Vector 13S-A (Addgene #48323). Point mutations and deletions were generated using PCR mutagenesis. For expression of DNAP^N+C^, vectors encoding His_6_-gp82 and untagged gp50 were co-transformed into *E. coli* Rosetta2 pLysS cells (EMD Millipore), then 1 L cultures in 2 L flasks were grown at 37°C to an A_600_ of 0.8, induced with 0.3 mM IPTG, then grown 16-18 hours at 20°C. Cells were harvested by centrifugation and resuspended in Buffer A (20 mM Tris-HCl pH 7.5, 300 mM NaCl, 10% glycerol, and 2 mM β-mercaptoethanol) plus 10 mM imidazole. Cells were lysed by sonication in the presence of protease inhibitors (Aprotinin, Pepstatin, Leupeptin, PMSF) and DNase I. Cell debris was pelleted by centrifugation, and proteins were passed over a Ni^2+^ affinity chromatography column (Ni-NTA Superflow, Qiagen), washed with Buffer A plus 20 mM imidazole, and eluted with Buffer A plus 400 mM imidazole. Eluted protein was concentrated and then passed over a size exclusion column (Superdex 200 Increase 10/300 GL, Cytiva) in SEC Buffer (20 mM Tris-HCl pH 7.5, 250 mM NaCl, 2 mM β-mercaptoethanol), and protein-containing fractions were concentrated and stored at 4°C. For expression of His_6_-Rhc, the same expression and purification procedure was followed.

For SYNZIP fusions, PCR-based mutagenesis was used to insert a C-terminal linker plus SYNZIP1 peptide (sequence below) to the untagged gp50 expression vector, and a C-terminal linker plus SYNZIP2 peptide (sequence below) to the His_6_-tagged gp155 expression vector. SYNZIP fusion proteins were expressed and purified as above.

<u>Linker</u> + SYNZIP1 sequence: <u>GSGSG</u>NLVAQLENEVASLENENETLKKKNLHKKDLIAYLEKEIANLRKKIEE

<u>Linker</u> + SYNZIP2 sequence: <u>GSGSG</u>ARNAYLRKKIARLKKDNLQLERDEQNLEKIIANLRDEIARLENEVASHEQ

For size exclusion chromatography linked to multi-angle light scattering (SEC-MALS), 100 µL of purified protein at 5 mg/mL was passed over a size exclusion column (Superdex 200 Increase 10/300 GL, Cytiva) in SEC buffer, followed by light scattering and differential refractive index (dRI) detectors (Wyatt Technology miniDAWN TREOS and Optilab T-rEX). Data were processed by ASTRA version 8 (Wyatt Technology).

### Single-particle cryoEM sample preparation and data collection

For PhiKZ DNAP^N+C^, purified protein was diluted to 10 µM in SEC buffer. 3 µL samples were applied to glow-discharged 300-mesh UltrAufoil grids (Electron Microscopy Sciences) in a Vitrobot Mark IV maintained at 4°C and 95% humidity (Thermo Fisher Scientific), blotted for 4 seconds with a blot force of 4.0, then vitrified in liquid ethane cooled by liquid nitrogen. The sample was imaged using a FEI Titan Krios G4 transmission cryoelectron microscope with Selectris X energy filter and Falcon 4 direct electron detector (Thermo Fisher Scientific) operating at 300 keV. The microscope was operated in EFTEM mode with a slit-width of 20 eV and using a 100 μm objective aperture. Automated data acquisition was performed using EPU (Thermo Fisher Scientific). Ten-second movies were collected at a magnification of 130,000x and a pixel size of 0.935 Å, with a total dose of 55 e-/Å^2^ distributed uniformly over 40 frames. In total, 4,222 movies were acquired with a realized defocus range of -1.2 to -2.0 μm. After curation of micrographs based on ice thickness and CTF fitting, 3,722 were accepted for particle picking.

All data processing (**Figure S1B-G**) was performed in cryoSPARC version 4 (Structura Biotechnology). For data processing, initial particle picking using the blob picker followed by 2D classification resulted in 2D classes used for template-based picking of an initial set of 1.8 million particles. 2D classification, *ab initio* model reconstruction, and non-uniform (NU) refinement resulted in a 2.4 Å resolution map calculated from 659,170 particles. This map was used to generate templates for a second round of template-based particle picking, resulting in a set of 3.6 million particles. Several rounds of 2D classification yielded a set of 1.4 million particles, which were subjected to heterogeneous refinement with three input 3D classes (which included the prior 2.4 Å reconstruction and a “junk” class), followed by NU refinement on a final set of 1,168,459 particles, yielding a 2.14 Å resolution map (**Table S1**).

For PhiKZ DNAP^N+C^–SYNZIP1 plus Rhc–SYNZIP2, separately purified proteins were mixed at 1:1 molar ratio, incubated 60 minutes at 4°C, then passed over a size exclusion column (Superdex 200 Increase, Cytiva) (**Figure S7A-B**). Fractions showing equimolar amounts of the three proteins were pooled, concentrated to 8.5 µM, and 3 µL samples were applied to glow-discharged 300-mesh UltrAufoil grids (Electron Microscopy Sciences) in a Vitrobot Mark IV maintained at 4°C and 95% humidity (Thermo Fisher Scientific), blotted for 4 seconds with a blot force of 4.0, then vitrified in liquid ethane cooled by liquid nitrogen. The sample was imaged using a FEI Titan Krios G4 transmission cryoelectron microscope with Selectris X energy filter and Falcon 4 direct electron detector (Thermo Fisher Scientific) operating at 300 keV. The microscope was operated in EFTEM mode with a slit-width of 20 eV and using a 100 μm objective aperture. Automated data acquisition was performed using EPU (Thermo Fisher Scientific). Ten-second movies were collected at a magnification of 105,000x and a pixel size of 0.83 Å, with a total dose of 50 e-/Å^2^ distributed uniformly over 40 frames. In total, 7,866 movies were acquired with a realized defocus range of - 1.2 to -2.0 μm. After curation of micrographs based on ice thickness and CTF fitting, 5,658 were accepted for particle picking.

For data processing (**Figure S7C-G**), initial particle picking using the blob picker followed by 2D classification resulted in a particle set of 24,693 particles which were subsequently subjected to *ab initio* reconstruction (2 classes) followed by NU refinement to yield a 3.73 Å map (17,131 particles). This map was used to generate templates for a second round of template-based particle picking, resulting in a set of 3.1 million particles. Several rounds of 2D classification yielded a final set of 184,433 particles, which were subjected to NU refinement, yielding a 2.82 Å resolution map (**Table S1**).

For the DNA-bound For PhiKZ DNAP^N+C^–SYNZIP1 plus Rhc–SYNZIP2 complex, proteins were reconstituted as above, and a primer-template DNA was assembled by annealing a 50-base template strand (5′-CGGTACCCGGGGATCCTCTAGAGTCGACCTGCAGGCATGCAAGCTTGGCA-3′) with a 16-base primer (5′-TGCCAAGCTTGCATGC-3′) with the 3′ cytosine carrying a 2′,3′-dideoxyribose to prevent catalysis. The cryoEM sample was prepared by mixing protein (final concentration: 8 µM) with 5x molar excess of DNA (final concentration: 40 µM) in a buffer containing 20 mM Tris-HCl pH 8.0, 10 mM MgCl_2_, 118 mM NaCl, 1 mM β-mercaptoethanol, and 0.5 mM dCTP, then incubating at room temperature for 1 hour. Following incubation, 3.2 µL of the combined sample was applied to glow-discharged 300-mesh UltrAufoil grids (Electron Microscopy Sciences) in a Vitrobot Mark IV maintained at 4°C and 95% humidity (Thermo Fisher Scientific), blotted for 4 seconds with a blot force of 4.0, then vitrified in liquid ethane cooled by liquid nitrogen and imaged on a Titan Krios microscope (Thermo Fisher Scientific) equipped with a K3 direct electron detector (Gatan) and a Biocontinuum Energy Filter (Gatan) operating at 300 keV. The microscope was operated in EFTEM mode with a slit-width of 20 eV and using a 100 μm objective aperture. Automated data acquisition was performed using EPU (Thermo Fisher Scientific). Ten-second movies were collected at a magnification of 105,000x and a pixel size of 0.83 Å, with a total dose of 50 e-/Å^2^ distributed uniformly over 40 frames. In total, 5,188 movies were acquired with a realized defocus range of -0.5 to -2.5 μm. After curation of micrographs based on ice thickness and CTF fitting, 4,680 were accepted for particle picking.

For data processing (**Figure S8**), initial particle picking was performed in cryoSPARC Live using the template picker, with templates generated from the final refined EM map of the PhiKZ DNAP^N+C^–SYNZIP1 plus Rhc–SYNZIP2 complex. Particles were selected from streaming 2D classification and separate manual iterative 2D classification, then combined and duplicate particles removed to give a set of 353,249 particles. These were subjected to *ab initio* reconstruction (2 classes) followed by heterogeneous refinement (one junk class and one protein class), resulting in a set of 256,596 particles. These were subjected to NU refinement, resulting in a 2.93 Å resolution map with interpretable density for a double-stranded DNA segment bound between the Rhc clamp domain and the DNAP thumb domain (**Figure S8A**). 3D classification was performed using a mask covering the bound DNA, resulting in two classes: one with no DNA bound, and one with DNA bound. Re-extraction, Global CTF refinement, and NU refinement on the final 137,984 DNA-bound particles resulted in a final 2.76 Å resolution map (**Table S1**).

Initial models of DNAP^N+C^, Rhc, and DNA were generated using AlphaFold 3 ^34^. Models were docked into cryoEM maps using ChimeraX^35^, followed by manual fitting and model rebuilding in Coot ^36^ and real-space refinement in phenix.refine^37^ (**Table S1**). Structural homologs were identified using Foldseek^20^ and DALI^19^ and structural comparison and analyses were performed using ChimeraX.

### *In vitro* polymerase and exonuclease assays

DNA polymerase and exonuclease activity assays were performed as previously described ^38^. Briefly, a template DNA strand was annealed to a 5′-6-FAM labelled DNA primer (see below for sequences) by mixing equimolar quantities in annealing buffer (10 mM Tris, pH 7.5, 50 mM NaCl, 1 mM EDTA), heating to 95°C for 5 minutes, then slow-cooled over 30 minutes in a PCR machine. All reactions were performed with 25 nM annealed template-primer, 300 nM PhiKZ DNAP^N+C^ (wild type, exonuclease-dead (gp82 D151A/E153A), polymerase-dead (gp82 D562A), or exonuclease+polymerase-dead), and 200 µM each dNTP in polymerase assay buffer (50 mM Tris, pH 7.5, 1 mM DTT, 2 mM MgCl_2_). 10 µL reactions were incubated at 25°C for varying timepoints, then reactions were stopped by the addition of 10ul of TBE-Urea Sample Buffer (Novex) supplemented with 10mM EDTA, followed by separation on a 15% Urea-PAGE gel and visualizing the bands with the Cy2 channel (492 nm excitation) on a Bio-Rad Chemidoc MP imager. For reactions with Rhc, 300 nM separately-purified Rhc was included. DNA sequences used: Template: 5′- CGGTACCCGGGGATCCTCTAGAGTCGACCTGCAGGCATGCAAGCTTGGCA-3′; Primer: 5′-(6-FAM)- TGCCAAGCTTGCATGCC-3′.

## DATA AVAILABILITY

Refined cryoEM maps and model coordinates have been deposited at the Electron Microscopy Data Bank (https://www.ebi.ac.uk/emdb/) and RCSB Protein Data Bank (https://www.rcsb.org), respectively, with the following accession codes:

DNAP^N+C^: EMDB EMD-48917; PDB 9N52
DNAP^N+C^-Rhc: EMDB EMD-78633; PDB 37XZ
DNAP^N+C^-Rhc + DNA: EMDB EMD-79225; PDB 38YB

## ACKNOWLEDGEMENTS

The authors thank members of the Pogliano, Villa, and Corbett labs for helpful conversations. The authors acknowledge support from the HHMI Emerging Pathogens Initiative grant (to E.V., J.P., and K.D.C.); National Institutes of Health (R01 GM129245 to J.P., R35 GM144121 to K.D.C.) and National Science Foundation MRI grant NSF DBI 1920374 (to E.V.); E.V. is a Howard Hughes Medical Institute Investigator. The authors acknowledge the use of the UC San Diego cryo-EM facility, part of the Goeddel Family Technology Sandbox, along with the scientific and technical assistance of facility staff members Dr. Mariusz Matyszewski and Dr. Inga Kuschnerus.

## AUTHOR CONTRIBUTIONS

D.B. determined cryoEM structures, performed biochemical assays, prepared figures, and wrote the manuscript with K.D.C. R.K.M. performed protein purification and performed biochemical assays. K.I. performed AlphaFold predictions and data analysis. J.P. provided advice on experimental design and data interpretation, and edited the manuscript. E.V. provided advice on experimental design and data interpretation, and edited the manuscript. K.D.C. oversaw the project, performed data analysis, prepared figures, and wrote the manuscript with D.B.

## SUPPLEMENTAL INFORMATION

**Table S1.** Cryoelectron microscopy data collection and refinement.

|  | PhiKZ DNAP<sup>N+C</sup> | PhiKZ DNAP<sup>N+C</sup>-Rhc | PhiKZ DNAP<sup>N+C</sup>-Rhc + DNA |
| --- | --- | --- | --- |
| **Data collection and processing** |  |  |  |
| Magnification | 130,000 | 105,000 | 105,000 |
| Voltage (kV) | 300 | 300 | 300 |
| Electron exposure (e-/Å<sup>2</sup>) | 55 | 50 | 50 |
| Defocus range (μm) | -1.2 to -2.0 | -1.2 to -2.0 | -0.5 to -2.5 |
| Pixel size (Å) | 0.935 | 0.83 | 0.83 |
| Symmetry imposed | none | none | none |
| Initial particle images (no.) | 3,621,040 | 3,130,367 | 3,792,792 |
| Final particle images (no.) | 1,168,459 | 184,433 | 137,984 |
| Map resolution (Å)<br>FSC threshold | 2.14 (0.143) | 2.82 (0.143) | 2.76 (0.143) |
| **Refinement** |  |  |  |
| Model resolution (Å)<br>FSC threshold | 2.1 (0.143) | 2.8(0.143) | 2.7(0.143) |
| Model composition |  |  |  |
| Non-hydrogen atoms | 11247 | 13068 | 13708 |
| Protein residues | 1340 | 1621 | 1620 |
| Water | 405 | 0 | 0 |
| Ligands | 1 Mg<sup>2+</sup> | 0 | 0 |
| *B* factors (Å<sup>2</sup>) |  |  |  |
| Protein | 69.98 | 142.38 | 109.59 |
| Water | 64.27 | N/A | N/A |
| Ligand | 68.03 | N/A | N/A |
| R.m.s. deviations |  |  |  |
| Bond lengths (Å) | 0.003 | 0.003 | 0.003 |
| Bond angles (°) | 0.446 | 0.523 | 0.559 |
| Validation |  |  |  |
| MolProbity score | 1.20 | 1.36 | 2.2 |
| Clashscore | 2.96 | 6.48 | 11.60 |
| Poor rotamers (%) | 1.42 | 0.9 | 3.12 |
| Ramachandran plot |  |  |  |
| Favored (%) | 98.5 | 98.08 | 96.34 |
| Allowed (%) | 1.5 | 1.92 | 3.66 |
| Disallowed (%) | 0 | 0 | 0 |
| PDB ID | 9N52 | 37XZ | 38YB |
| EMDB ID | EMD-48917 | EMD-78633 | EMD-79225 |

**Table S2.** PhiKZ Rhc C-terminal domain DALI hits. Top 25 hits from the PDB25 subset. Light green highlight indicates sliding clamp proteins

| PDB-Chain | Z score | Calpha rmsd | Alignment length | Uniprot ID | Description |
| --- | --- | --- | --- | --- | --- |
| 3ggr-B | 6.3 | 4.6 | 179 | O60921 | *H. sapiens* HUS1 |
| 7st9-G | 5.7 | 3.9 | 165 | Q08949 | *S. cerevisiae* Ddc1 |
| 3a1j-A | 5.6 | 3.7 | 154 | Q99638 | *H. sapiens* RAD9A |
| 7ype-A | 5.5 | 4 | 155 | A0A0A1E3R5 | African swine fever virus E301R |
| 7st9-H | 5.4 | 4.7 | 167 | Q02574 | *S. cerevisiae* Mec3 |
| 3ggr-C | 4.9 | 4.1 | 155 | O60671 | *H. sapiens* RAD1 |
| 6drt-A | 4.6 | 4 | 148 | P04525 | phage T4 gp45 |
| 4h5j-B | 4.5 | 14.7 | 59 | P11655 | *S. cerevisiae* SEC12 |
| 3elz-B | 4.4 | 9.5 | 64 | Q6IMW5 | *Danio rerio* ileal bile acid binding protein |
| 3cnx-C | 4.4 | 3.8 | 60 | Q82EE4 | *Streptomyces avermitilis* putative dehydratase |
| 3h3h-B | 4.4 | 4.2 | 57 | Q2T8S1 | *Burkholderia thailandensis* E264 SnoaL-like protein |
| 2awa-B | 4.3 | 4.2 | 155 | O06672 | *Streptococcus pneumoniae* TIGR4 (DNA Pol III beta chain) |
| 4hvt-A | 4.3 | 17.9 | 67 | Q68XJ3 | *Rickettsia typhi* post-proline cleaving enzyme |
| 3vpd-A | 4.3 | 2.6 | 79 | Q5SH23 | *T. thermophilus* LysX |
| 6qp8-A | 4.3 | 8.8 | 69 | A0A0B4KG38 | *D. melanogaster* semaphorin 2b |
| 6lk8-J | 4.3 | 13.2 | 80 | A0A1L8H1I9 | *X. laevis* Nup133 |
| 5mwv-A | 4.2 | 8.5 | 82 | P76045 | *E. coli* OmpG |
| 8cli-B | 4.2 | 11.4 | 68 | Q9UKN8 | *H. sapiens* TFIIIC polypeptide 4 |
| 4pr7-A | 4.2 | 8.7 | 59 | Q934G3 | *Dickeya dadantii* KdgM porin |
| 7uic-p | 4.2 | 16.3 | 71 | P32259 | *S. cerevisiae* Mediator subunit 16 |
| 6bjt-B | 4.2 | 4.1 | 61 | A0A1A5DB13 | *Pseudomonas* sp. EGD-AKN5 AtzH |
| 5nqz-A | 4.1 | 5.2 | 60 | Q9JXV4 | *Neisseria meningitidis* MC58 outer membrane protein |
| 1k32-A | 4.1 | 3.3 | 49 | P96086 | *Thermoplasma acidophilum* tricorn protease |
| 8rf0-A | 4.1 | 15 | 64 |  | *Agrobacterium tumefaciens* NdvB |
| 9gcz-A | 4 | 12 | 63 | Q8A622 | *Bacteroides thetaiotaomicron* VPI-5482 XusB |

**Figure S1.**
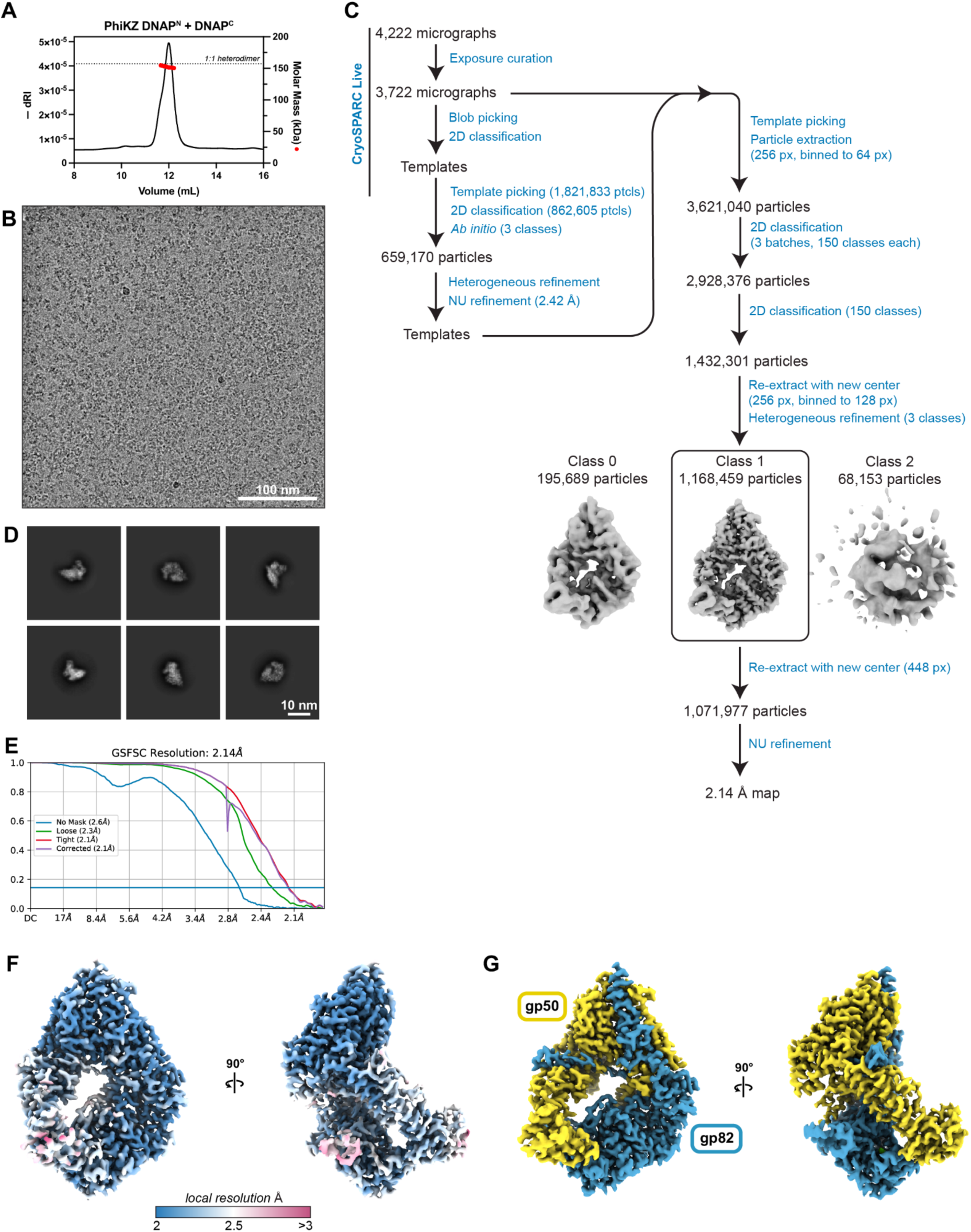
CryoEM structure determination of PhiKZ DNAP^N+C^. (A) Size exclusion chromatography coupled to multi-angle light scattering (SEC-MALS) analysis of purified PhiKZ DNAP^N+C^. Measured molecular weight (red circles): 152 kDa. Dashed line indicates expected molecular weight of a 1:1 heterodimer of gp82 and gp50: 157.2 kDa. dRI: differential refractive index; measurement of protein concentration (black line). (B) Example micrograph. Scale bar: 100 nm. (C) Data processing workflow. (D) Example 2D classes from 1.4 M-particle set used for 3D structure determination. (E) Fourier Shell Correlation (FSC) graph from final 2.14 Å resolution refinement. (F) Two views of the final 2.14 Å resolution refinement map, colored by local resolution. (G) Two views of the final 2.14 Å resolution refinement map, colored by subunit (DNAP^N^ (gp82) blue, DNAP^C^ (gp50) yellow).

**Figure S2.**
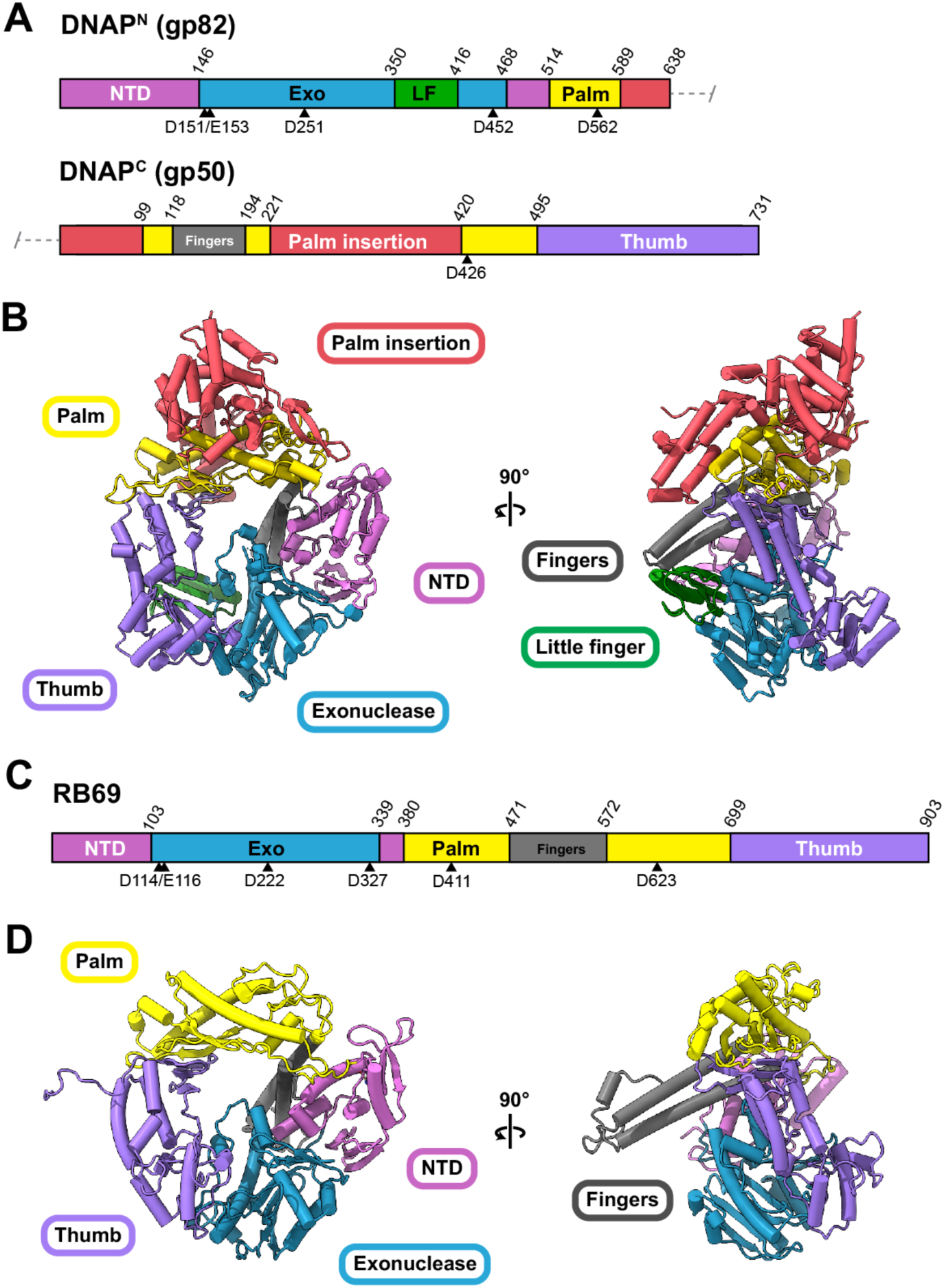
Comparison of PhiKZ and RB69 DNAP structures. (A) Domain diagram of the two-subunit PhiKZ DNA polymerase. (B) Two views of the PhiKZ DNAP^N+C^ complex, colored by domain as in panel (A). (C) Domain diagram of the RB69 DNA polymerase. (D) Structure of the RB69 DNAP (PDB ID 1WAJ)^15^, colored by domain as in panel (C).

**Figure S3.**
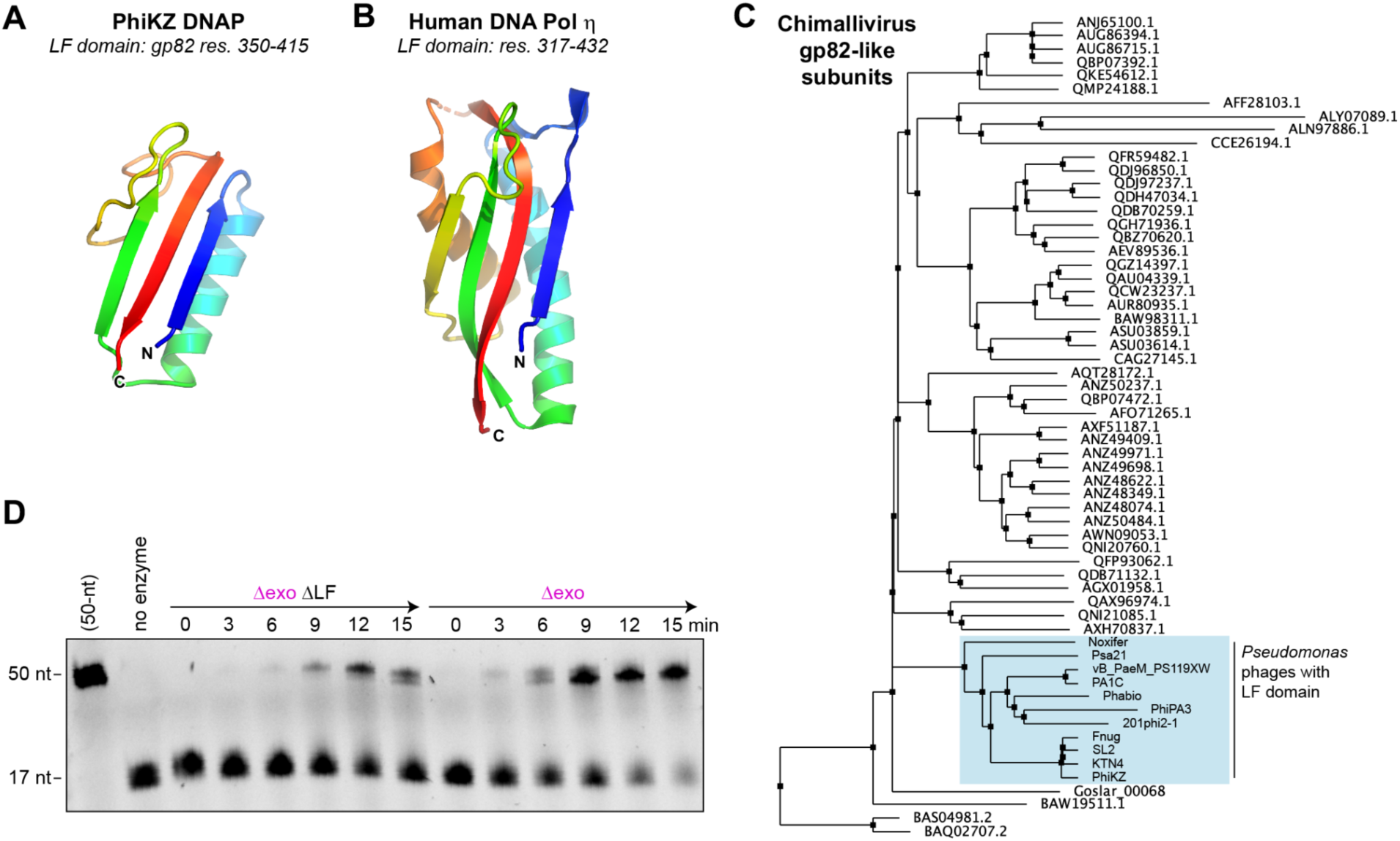
Structure of the PhiKZ DNAP LF domain. (A) Cartoon view of the little finger (LF) domain from PhiKZ DNAP^N^ (gp82 residues 350-415), colored as a rainbow from blue (N-terminus of the domain) to red (C-terminus of the domain). (B) Cartoon view of the LF domain of human DNA polymerase η (eta) (PDB ID 4J9S)^39^, aligned with the PhiKZ DNAP LF domain. The DALI server^19^ reports an r.m.s.d. of 3.0 Å over 66 Cɑ atoms when compared to the PhiKZ DNAP LF domain. (C) Neighbor-joining phylogenetic tree (calculated by JalView^40^) on a set of chimallivirus DNAP gp82-like subunits. All sequences are denoted by their NCBI accession numbers except for a set of *Pseudomonas* phages that possess the LF domain, labeled by their common names (shaded blue; NCBI accession numbers ARV77505 (Noxifer), QBJ02620 (Psa21), QEM41817 (vB_PaeM_PS119XW), QBX32238 (PA1C), ARV76759 (Phabio), AEH03509 (PhiPA3), ABY62974 (201phi2-1), QJB22744 (Fnug), ATN94847 (SL2), ANM44866 (KTN4), NP_803648 (PhiKZ)). (D) Primer extension assay on PhiKZ DNAP^N+C^ Δexo and a mutant missing the LF domain (PhiKZ DNAP^N+C^ Δexo ΔLF; gp82 residues 350-417 replaced by GSGA). For each enzyme, time points from 0 to 15 minutes are shown. Data are representative of three separate trials.

**Figure S4.**
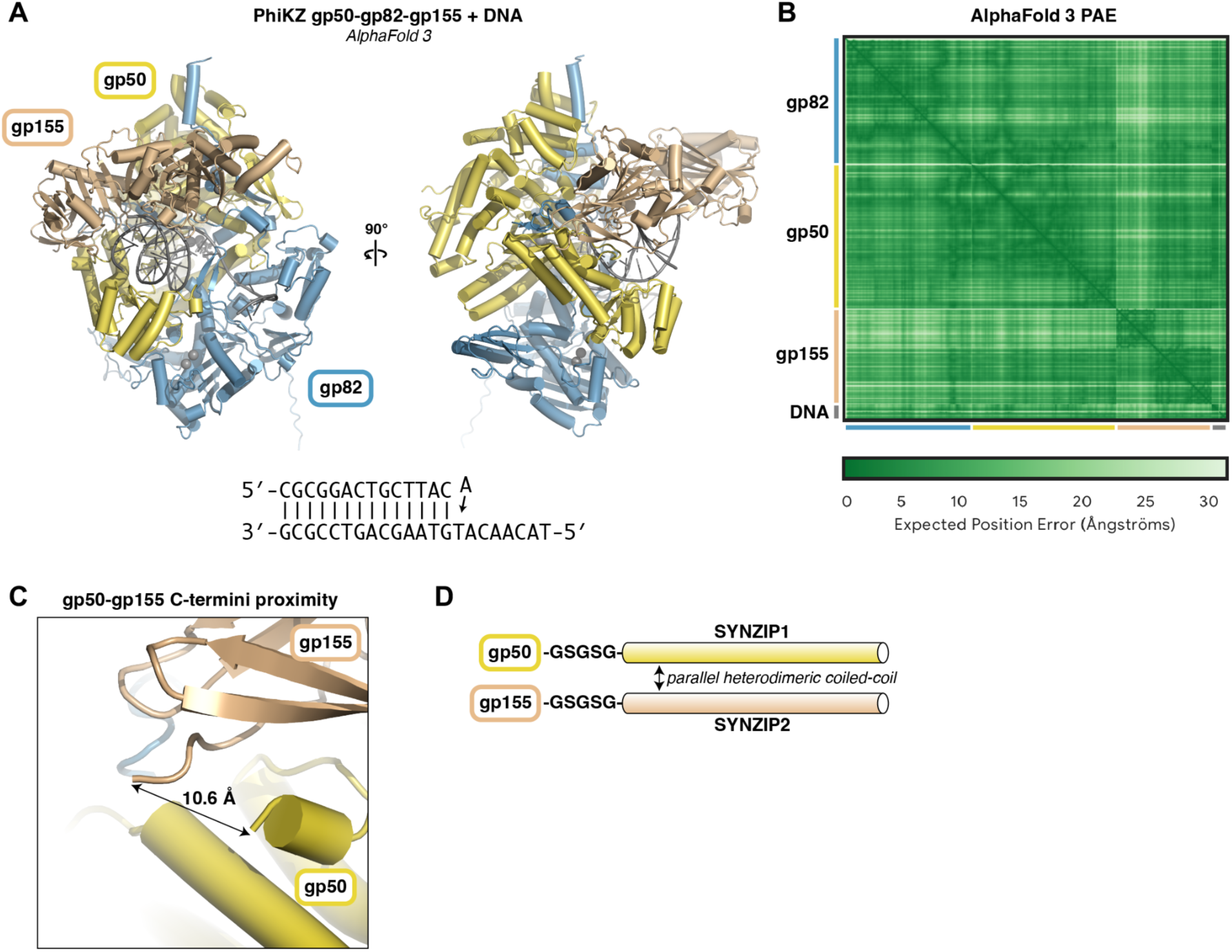
AlphaFold predicted structure of PhiKZ DNAP^N+C^-Rhc-DNA complex. (A) Two views of an AlphaFold 3 predicted structure of PhiKZ DNAP^N^ (gp82; blue), DNAP^C^ (gp50; yellow), and Rhc (gp155; brown) bound to a primer-template DNA (sequence at bottom), an incoming nucleotide (modeled as ATP), and four Mg2+ ions (two binding to the polymerase active site, two binding to the exonuclease active site). (B) AlphaFold 3 predicted aligned error (PAE) plot for the prediction shown in panel (A). (C) Closeup of the C-termini of gp50 and gp155, showing their close juxtaposition. (D) Schematic of the SYNZIP fusion strategy. SYNZIP1 and SYNZIP2 were previously shown to form a heterodimeric parallel coiled-coil ^28^. See **Materials and Methods** for SYNZIP sequences.

**Figure S5.**
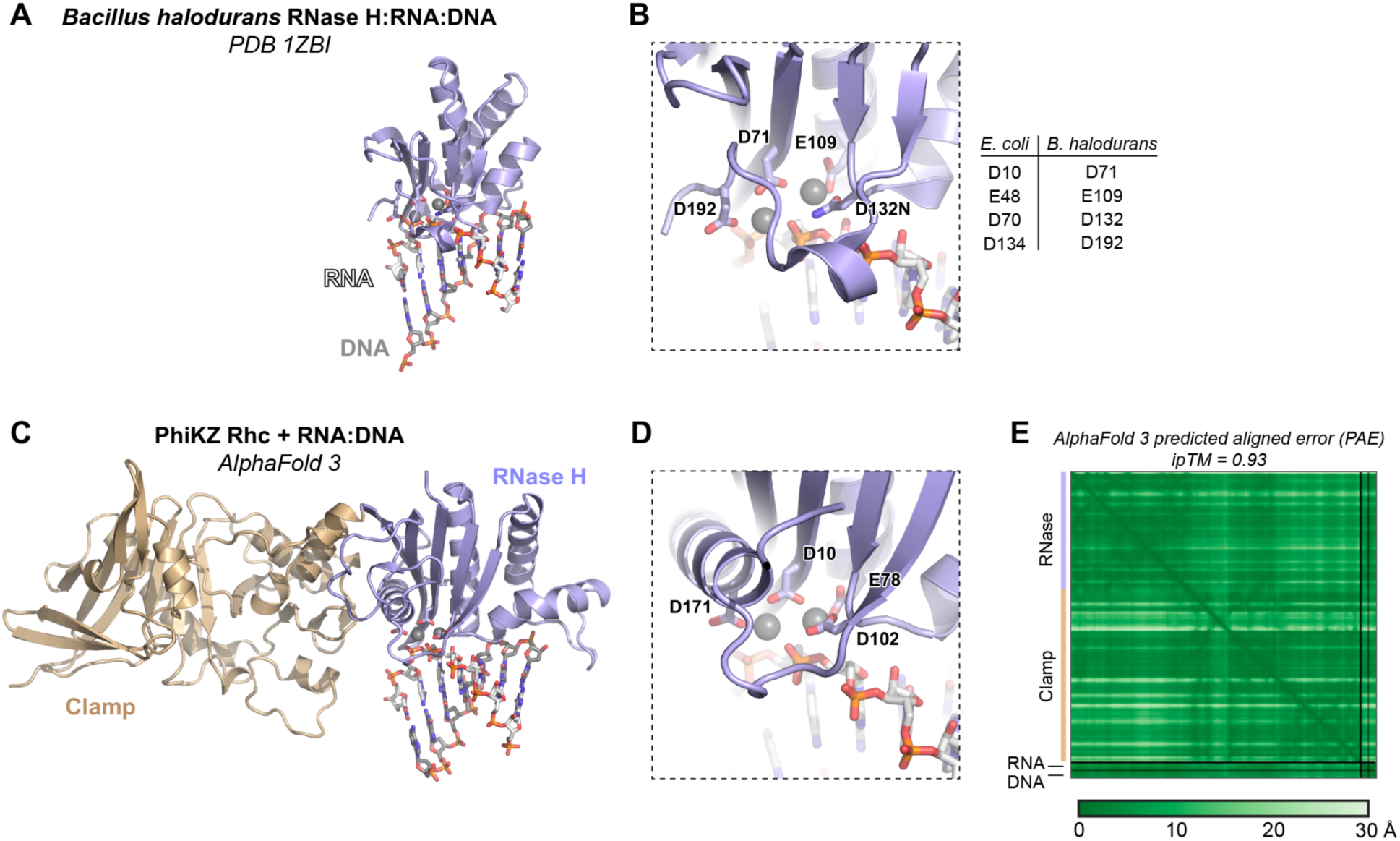
The Rhc N-terminal domain is related to RNase H1. (A) Structure of *B. halodurans* RNase H bound to an RNA-DNA hybrid (PDB ID 1ZBI) ^41^. (B) Closeup of the *B. halodurans* RNase H catalytic site, with conserved catalytic residues shown as sticks and two Mg^2+^ ions shown as spheres. *Right:* Equivalent active site residues of *E. coli* RNase H and *B. halodurans* RNase H. (C) AlphaFold 3 predicted structure of PhiKZ Rhc bound to an RNA:DNA hybrid and two Mg^2+^ ions. The N-terminal domain is colored blue; the C-terminal domain is colored light brown; RNA is colored white; DNA is colored gray; Mg^2+^ ions are shown as gray spheres. The N-terminal domain (blue) is shown in the same orientation as *B. halodurans* RNase H in panel (A). (D) Closeup of the predicted RNase H domain catalytic site of PhiKZ Rhc, with conserved catalytic residues shown as sticks and two Mg^2+^ ions shown as spheres. (E) AlphaFold 3 predicted aligned error (PAE) plot for the prediction shown in panel (C).

**Figure S6.**
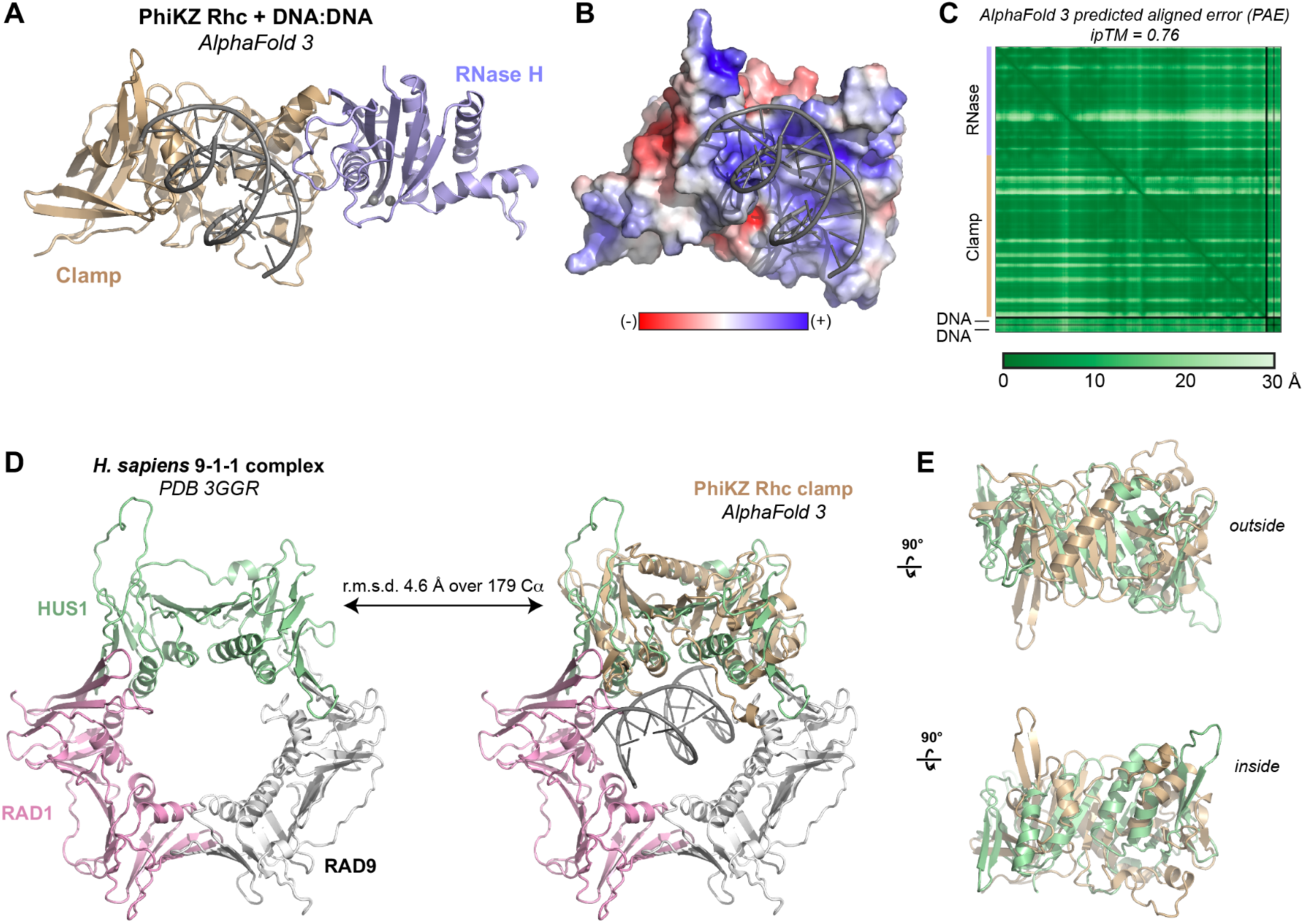
The Rhc C-terminal domain is related to DNA sliding clamps. (A) Predicted structure of PhiKZ Rhc bound to a double-stranded DNA. (B) Electrostatic surface of the PhiKZ Rhc C-terminal domain, colored by surface charge (red negative, blue positive). (C) AlphaFold 3 predicted aligned error (PAE) plot for the predicted structure of PhiKZ RNase H bound to a double-stranded DNA. (D) Structure of the *H. sapiens* 9-1-1 complex (PDB ID 3GGR)^42^, with the three subunits individually colored and labeled. *Center:* Overlay of the PhiKZ Rhc N-terminal domain predicted structure (with bound DNA) onto HUS1. (E) Two rotated views of the Rhc-HUS1 overlay.

**Figure S7.**
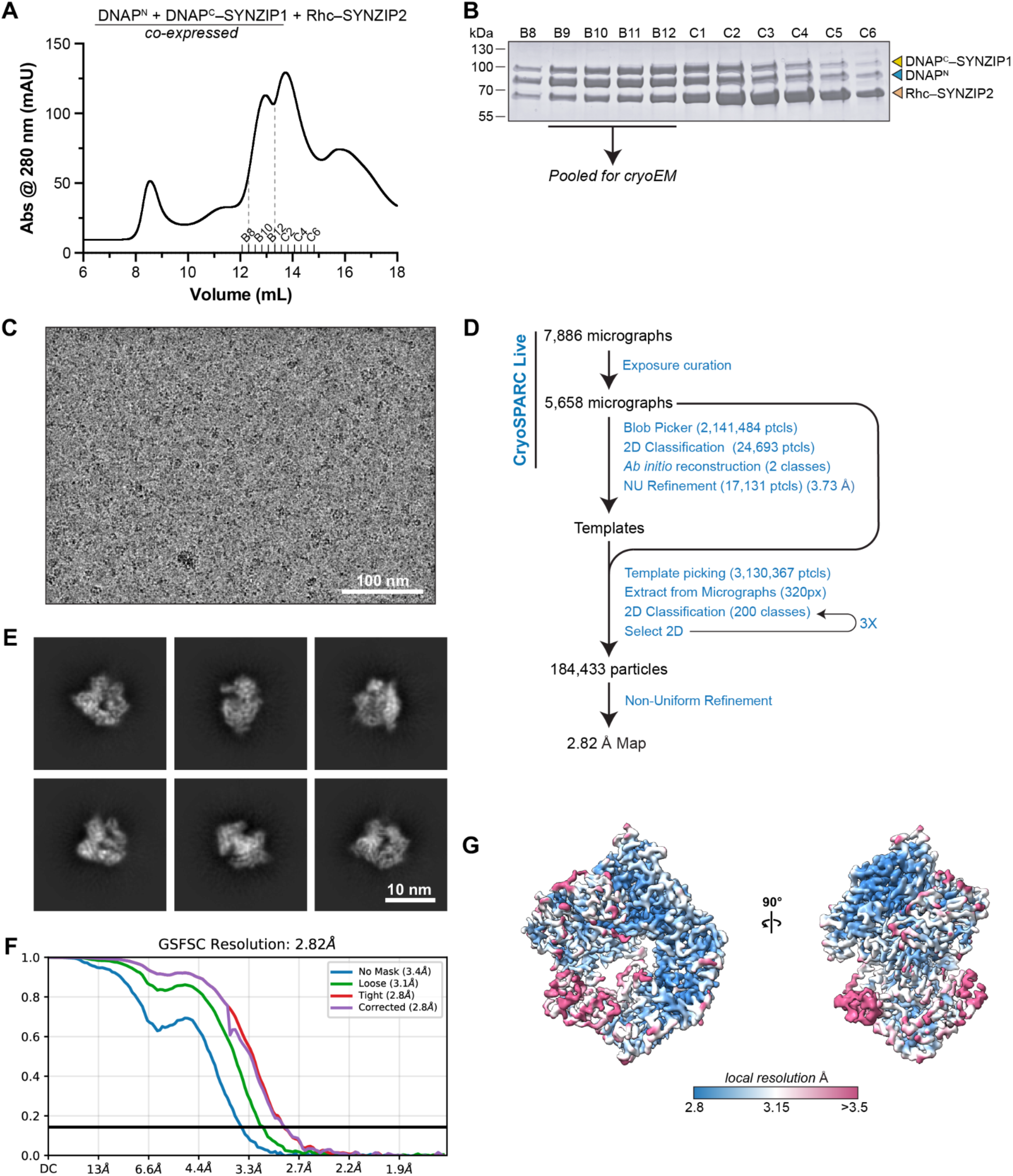
CryoEM structure determination of PhiKZ DNAP^N+C^ + Rhc. (A) Size exclusion chromatography (Superdex 200 Increase 10/300) for separately expressed (DNAP^N^ + DNAP^C^–SYNZIP1) and (Rhc–SYNZIP2), mixed at 1:1 molar ratio. Fraction labels correspond to lanes in panel (B), and dotted lines indicate fractions pooled for cryoEM. (B) SDS-PAGE analysis of size exclusion fractions from panel (A). Fractions B9-B12 were pooled for cryoEM. (C) Representative micrograph. (D) Structure determination workflow. (E) Representative 2D classes of the final 184,433 particle set used for refinement. (F) Fourier Shell Correlation (FSC) graph from final 2.82 Å resolution refinement. (G) Two views of the final 2.82 Å resolution refinement map, colored by local resolution.

**Figure S8.**
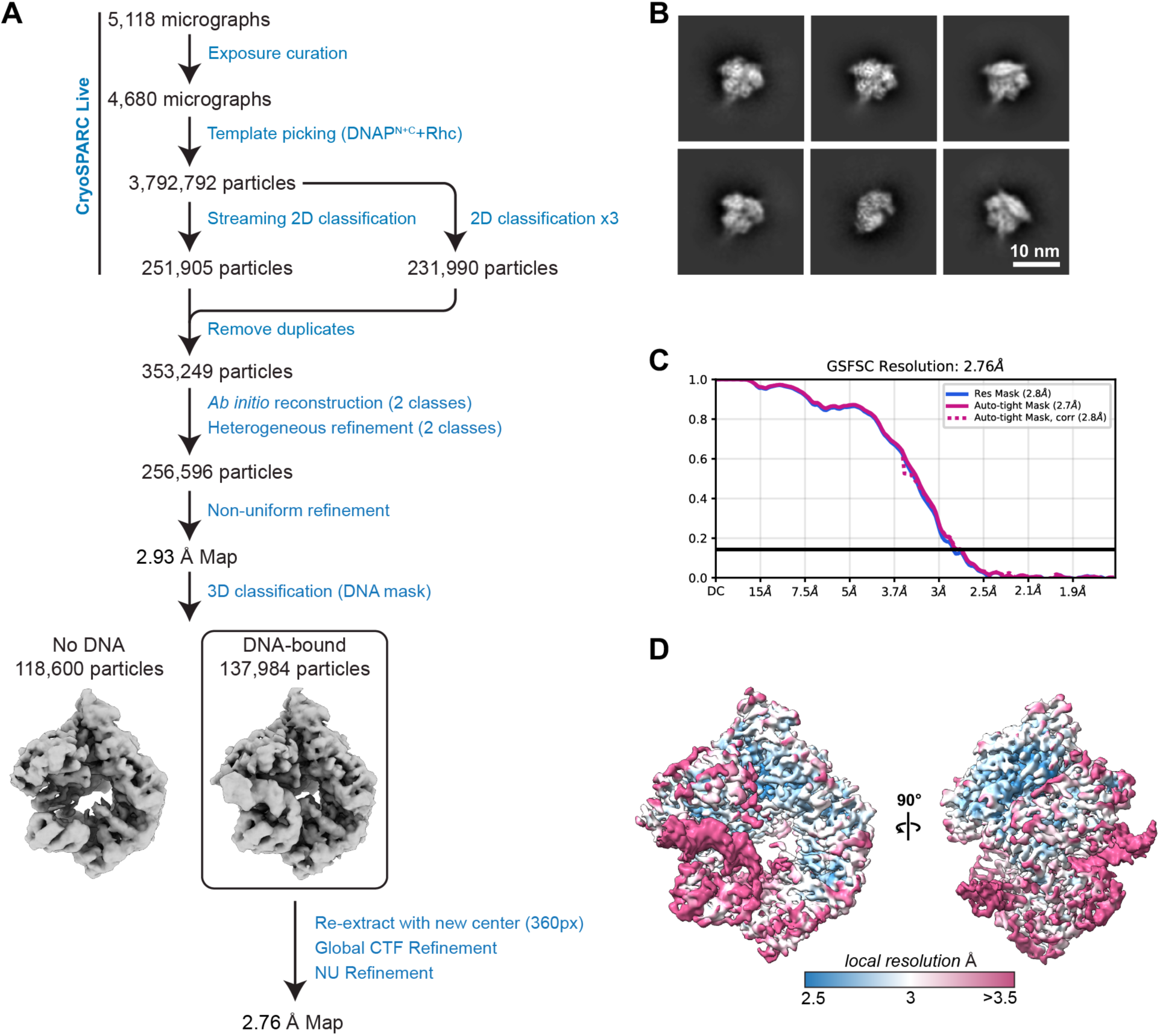
CryoEM structure determination of PhiKZ DNAP^N+C^ + Rhc + DNA. (A) Structure determination workflow. (B) Representative 2D classes of the final 137,984 particle set used for refinement. (C) Fourier Shell Correlation (FSC) graph from final 2.76 Å resolution refinement. (D) Two views of the final 2.76 Å resolution refinement map, colored by local resolution.

